# R2ucare: An R package to perform goodness-of-fit tests for capture-recapture models

**DOI:** 10.1101/192468

**Authors:** Olivier Gimenez, Jean-Dominique Lebreton, Rémi Choquet, Roger Pradel

## Abstract

1. Assessing the quality of fit of a statistical model to data is a necessary step for conducting safe inference.
2. We introduce `R2ucare`, an R package to perform goodness-of-fit tests for open single-and multi-state capture-recapture models. `R2ucare` also has various functions to manipulate capture-recapture data.
3. We remind the basics and provide guidelines to navigate towards testing the fit of capture-recapture models. We demonstrate the functionality of `R2ucare` through its application to real data.

## Introduction

Capture–recapture (CR) models have become a central tool in population ecology for estimating demographic parameters under imperfect detection of individuals (Lebreton et al. 1992; 2009). These methods rely on the longitudinal monitoring of individuals that are marked (or identifiable) and then captured or sighted alive over time.

Single-state CR models, and the Cormack-Jolly-Seber (CJS) model in particular (Lebreton et al. 1992), have been used to assess the effect of climate change (e.g. Guéry et al. 2017) or study senescence (e.g. Péron et al. 2016). The extension of single-state models to situations where individuals are detected in several geographical sites or equivalently states (e.g. breeding/non-breeding or sane/ill) are called multi-state CR models (Lebreton et al. 2009). Multistate CR models, and the Arnason-Schwarz model in particular (Lebreton et al. 2009), are appealing for addressing various biological questions such as metapopulation dynamics (e.g. Spendelow et al. 2016) or life-history trade–offs (e.g. Supp et al. 2015).

A necessary step for correct inference about demographic parameters is to assess the fit of single-and multi-state models to CR data, regardless of whether a Bayesian or a frequentist framework is adopted.

Two family of methods exist to perform goodness-of-fit (GOF) tests for CR models. First, an omnibus test of the null hypothesis that a given model fits the data adequately can be conducted using resampling methods and the deviance as a metric (White 2002). However when the null hypothesis is rejected, this omnibus approach does not inform about an alternative model that could be fitted. Second, specialized tests have been built to address biologically meaningful causes of departure from the null hypothesis. A global test for single-and multi-state CR models is decomposed into several interpretable components based on contingency tables, for example the presence of transients (Pradel et al., 1997; Pradel et al. 2003) or that for trap–dependence (Pradel, 1993; Pradel et al. 2003). These GOF tests are implemented in the Windows application U-CARE (Choquet et al. 2009).

Here, we introduce the R (R Development Core Team 2014) package `R2ucare` to perform GOF tests for single-and multi-state CR models. `R2ucare` also includes various functions to help manipulate CR data. As a package in the CRAN database, `R2ucare` provides full advantage of R’s many features (e.g. simulations, model fitting), while being multi-platform. We go through the theory first, then illustrate the use of `R2ucare` with an example on wolf in France for single-state models and geese in the U.S. for multi-state models.

## Theory

Once a model has been specified, GOF testing is the procedure that controls model assumptions. GOF testing and model fitting are two complementary procedures that share and compete for the information contained in the data. More liberal models require more information to be fitted (there are more parameters to estimate) but also fewer assumptions need to be verified. For instance, the time-dependent CJS model is merely content with the numbers of individuals captured at each occasion and the numbers never seen again from those released at each occasion when it comes to estimating its parameters. These summary statistics leave much of the details of the capture histories available to test its assumptions.

There are several ways in which this remaining information may be exploited to test the assumptions. The implementation retained in `R2ucare` builds on the optimal approach originally devised by Pollock et al. (1985) and later modified by Pradel (1993). It is based on contingency tables and aims at testing with chi-squared tests (and Fisher’s exact tests when needed) for transients and trap-dependence. These aspects are examined specifically in two independent component tests called respectively `Test 3.SR` and `Test 2.CT`. The component tests directed at transients and trap-dependence actually address features of the data that are consequences of respectively the presence of transients and trap-dependence, so that these features may also be caused by other, completely different phenomena. They do verify respectively that:

- Newly encountered individuals have the same chance to be later reobserved as recaptured (previously encountered) individuals; this is the null hypothesis of `Test 3.SR`.
- Missed individuals have the same chance to be recaptured at the next occasion as currently captured individuals; this is the null hypothesis of `Test 2.CT`.

Although these components are often called ‘test of transience’ and ‘test of trap-dependence’, when it comes to interpretation, one should keep in mind that transience and trap-dependence are just two specific reasons why the tests respectively called `3.SR` and `2.CT` might be significant.

Interestingly, these two components provide formal tests for comparing the CJS model with more general models, namely a model with an interaction between age (2 classes) and time in the survival probability for `Test 3.SR` (Pradel et al. 1997) and a model allowing for a different recapture probability of individuals just released for `Test 2.CT` (Pradel 1993).

Beyond these two oriented components, the remaining information is distributed and structured into two additional components: `Test 3.Sm` and `Test 2.CL`. Those examine long-term features of the data:

- Among those individuals seen again, when they were seen does not differ among previously and newly marked individuals; this is the null hypothesis of `Test 3.Sm`.
- There is no difference in the timing of reencounters between the individuals encountered and not encountered at occasion *i*, conditional on presence at both occasions *i* and *i* + 2; this is the null hypothesis of `Test 2.CL`?

Data are generally sparse for these components and scattered over many occasions. Despite the implementation of some automatic pooling (see Choquet et al. 2005 for more details about the pooling rules), they are rarely significant alone.

Although many situations can lead to similar test results, we propose here a decision tree (Figure 1) that should lead to reasonable solutions in most cases.

**Figure 1:**
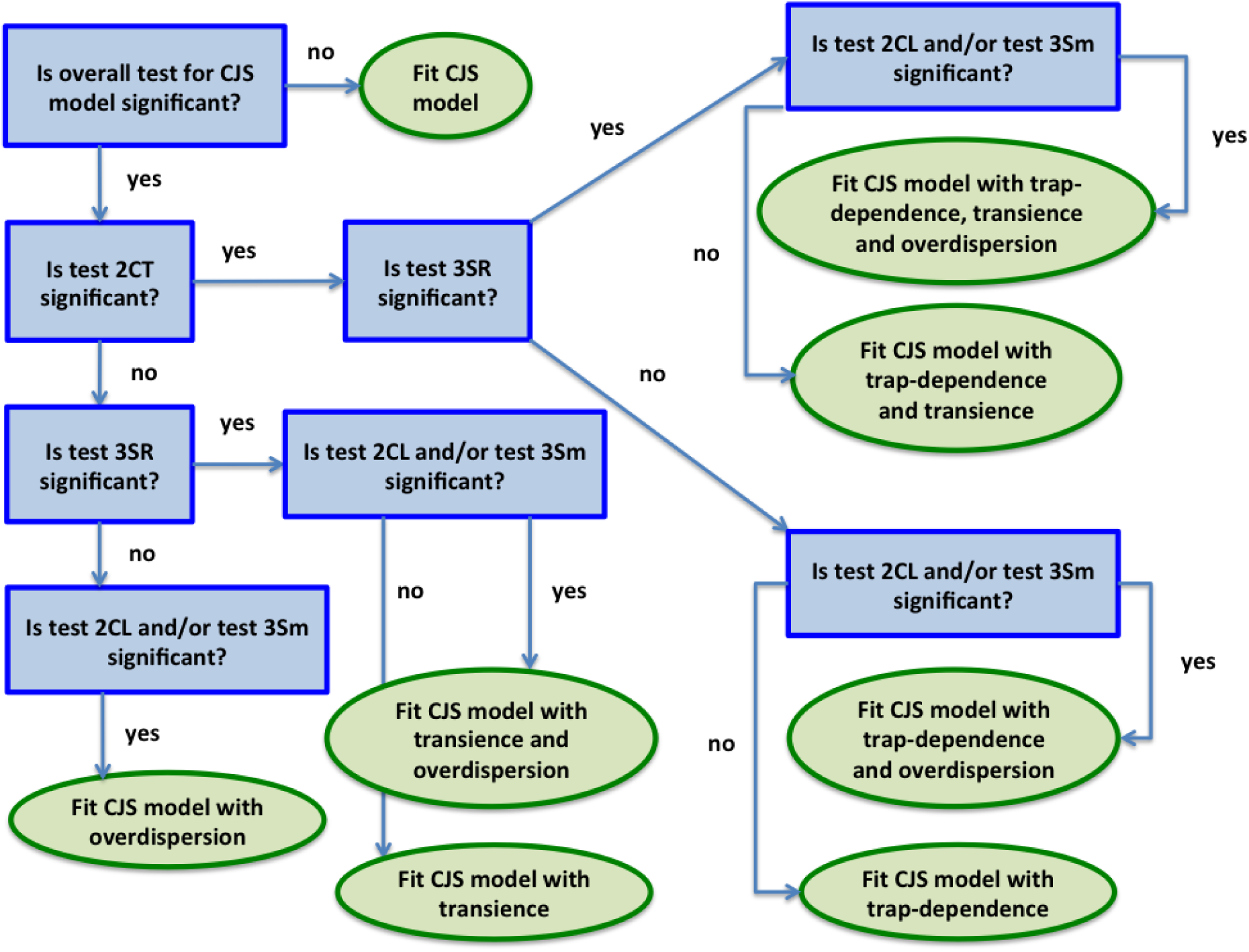
Decision tree to navigate towards testing the fit of single site/state capture-recapture models, with the Cormack-Jolly-Seber (CJS) model as a reference. Questions are in the blue rectangles, actions in the green ellipses. We start by asking the question in the top-left corner. The coefficient of overdispersion is calculated as the ratio of the goodness-of-fit test statistic over the number of degrees of freedom (Pradel et al. 2005). *Remark 1*: we begin by testing for the presence of trap-dependence, then that of transience; these steps could be permuted without affecting the final outcome. *Remark 2*: the overall goodness-of-fit test may be significant while none of the four sub-components is; in this situation, we recommend fitting the CJS model and correcting for overdispersion. *Remark 3*: we do not cover the issue of heterogeneity for which a formal test does not exist. When both the tests for the presence of transience and trap-dependence are significant, and only them, there is suspicion of heterogeneity in detection (Péron et al. 2010). Péron et al. (2010) implemented an approximate procedure to assess the presence of heterogeneity in the detection process, and Jeyam et al. (2017) developed a formal test for the same purpose. Cubaynes et al. (2012) recommended using the Akaike Information Criterion (AIC) to compare models with and without heterogeneity. *Remark 4*: To account for the presence of transience, that of trap-dependence or an effect of heterogeneity, we refer to Pradel et al. (1997), Pradel and Sanz-Aguilar (2012; see also Pradel 1993 and Gimenez et al. 2003) and Gimenez et al. (2017) respectively.

The theory for the GOF test of the multistate Arnason-Schwarz model was developed along similar lines as for the CJS model (Pradel et al. 2003). This test has yet more components and some components have a more complex structure (hence our non attempt to build a decision tree as for the CJS model), but for all that concerns us, the reasoning remains very similar. The test implemented in `R2ucare` is actually a test of the Jolly-Move model, a slightly more general model than the Arnason-Schwarz model in that it allows detection parameters to depend on the previous state occupied. This is biologically irrelevant in most common situations (Pradel et al. 2003), so that we may reason as if we were examining the Arnason-Schwarz model. Components here have been designed to detect transients, trap-dependence, and the memory of past states. This last point means that the component examines whether transitions to a new state depend on previous states beyond the current one. The corresponding components are respectively `Test 3.GSR`, `Test M.ITEC`, and `Test WBWA`. Like for the CJS case, they actually examine features of the data, namely that:

- Newly encountered individuals have the same chance to be later reobserved as recaptured (i.e. previously encountered) individuals; this is the null hypothesis of `Test 3.GSR` which is the exact equivalent of `3.SR`.
- Individuals currently in the same state, whether captured or missed, have the same chance to be recaptured in each state at the next occasion; this is the null hypothesis of `Test M.ITEC`.
- Individuals currently captured in the same state have the same chance to be next reobserved in the different states independently of their observed state at the most recent capture; this is the null hypothesis of `Test WBWA`.

These interpretable components are complemented by two composite components with no clearly identified interpretation, `Test 3.GSm` and `Test M.LTEC`. We do not attempt to give a description of these; let it suffice to say that `Test 3.GSm` is concerned with comparing newly and previously encountered, while `Test M.LTEC` contrasts missed and encountered individuals. Fortunately, these components play a secondary role as they are usually not significant alone.

For more details about the theory of GOF testing for CR models, we strongly encourage users to read Pradel et al. (2005) and Cooch and White (2006).

## The R2ucare package

The `R2ucare` package contains R functions to perform GOF tests for CR models as well as various functions to manipulate CR data (see Table 1 and the vignette of the package named `vignette_R2ucare`). It ensures reproducibility which was not possible with the `U-CARE` (Choquet et al. 2009) Windows standalone application. Besides, it can be used in combination with other R packages for fitting CR data like `RMark` (Laake 2013) or `marked` (Laake et al. 2013) or to carry out simulations to assess statistical power (e.g. Bromaghin et al. 2013; Fletcher et al. 2012).

**Table 1:**
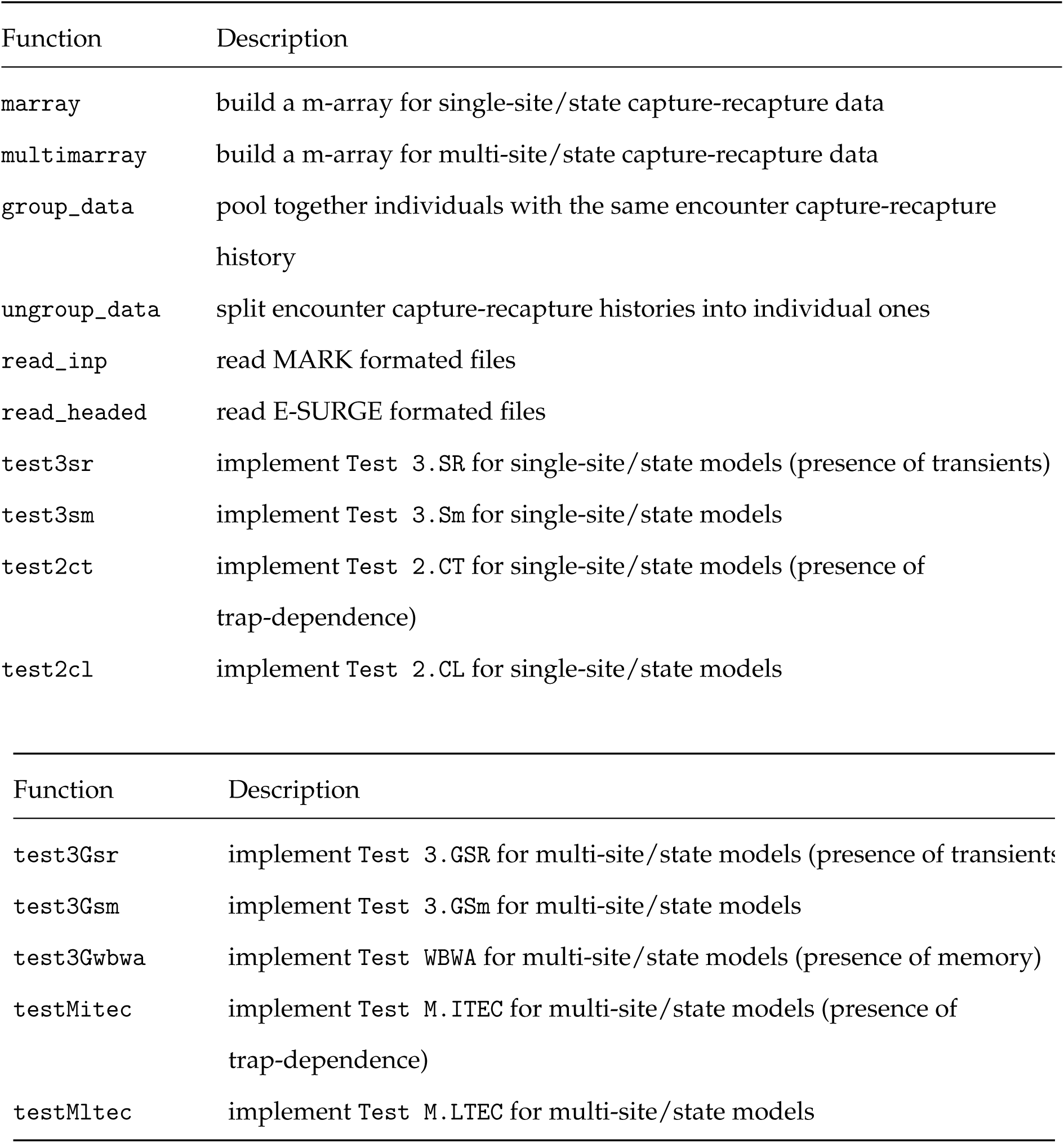
The main functions of `R2ucare` and their description. See main text for more details.

| Function | Description |
| --- | --- |
| marray | build a m-array for single-site/state capture-recapture data |
| multimarray | build a m-array for multi-site/state capture-recapture data |
| group_data | pool together individuals with the same encounter capture-recapture history |
| ungroup_data | split encounter capture-recapture histories into individual ones |
| read_inp | read MARK formatted files |
| read_headed | read E-SURGE formatted files |
| test3sr | implement Test 3.SR for single-site/state models (presence of transients) |
| test3sm | implement Test 3.Sm for single-site/state models |
| test2ct | implement Test 2.CT for single-site/state models (presence of trap-dependence) |
| test2cl | implement Test 2.CL for single-site/state models |
| test3Gsr | implement Test 3.GSR for multi-site/state models (presence of transients) |
| test3Gsm | implement Test 3.GSm for multi-site/state models |
| test3Gwbwa | implement Test WBWA for multi-site/state models (presence of memory) |
| testMitec | implement Test M.ITEC for multi-site/state models (presence of trap-dependence) |
| testMltec | implement Test M.LTEC for multi-site/state models |

## Goodness-of-fit tests for single-site/state models

We illustrate the use of `R2ucare` to assess the GOF of the CJS model to a dataset on wolves (*Canis lupus*) in France (e.g., Fletcher et al. 2012). Briefly, the data consist of capture histories for 160 individuals, partitioned into 35 3-month intervals (from spring 1995 to autumn 2003).

We first load the `R2ucare` package:

~~~
library(R2ucare)
~~~

Then we read in the wolf data that is provided with the package. To do so, `R2ucare` contains two functions that accomodate the most frequent CR formats: `read_inp` deals with the `MARK` format (Cooch and White 2006) while `read_headed` deals with the `E-SURGE` format (Choquet et al. 2009). The wolf dataset has the `MARK` format, therefore:

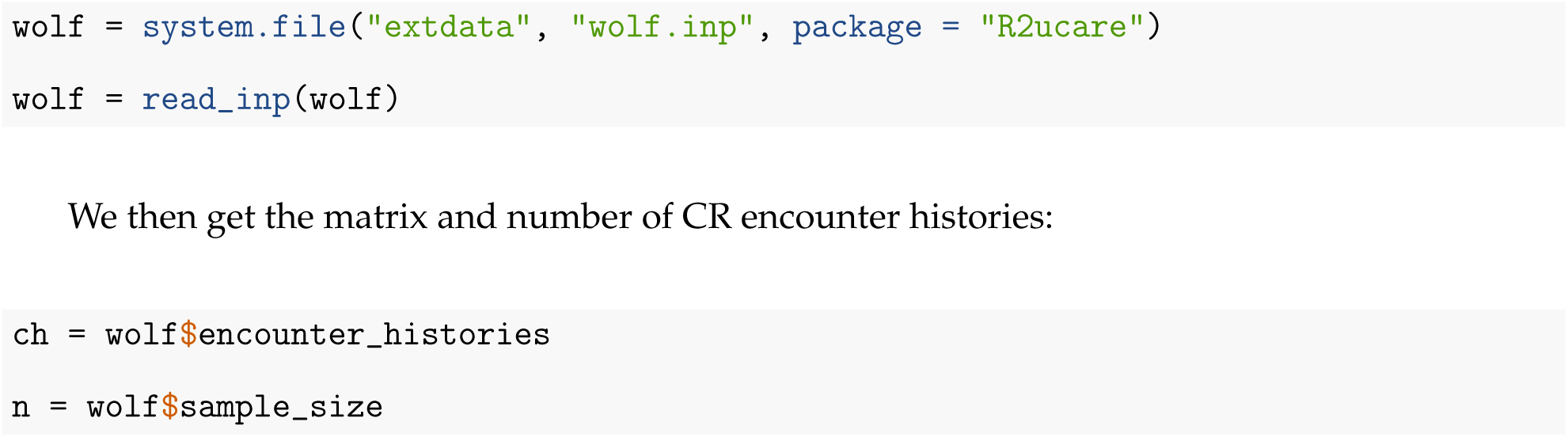

Following the procedure described in Figure 1, we first assess the overall fit of the CJS model by using the function `overall_CJS`:

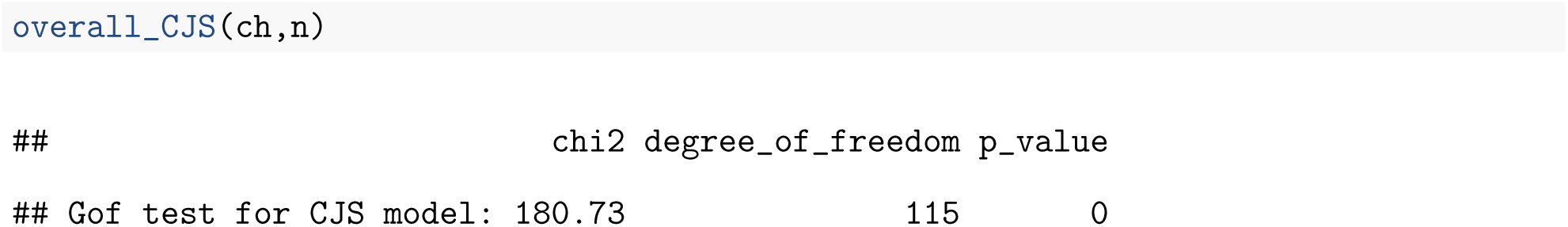

Clearly, the CJS model does not fit the data well (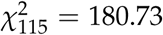, *P* < 0.01). We then test for an effect of trap-dependence:

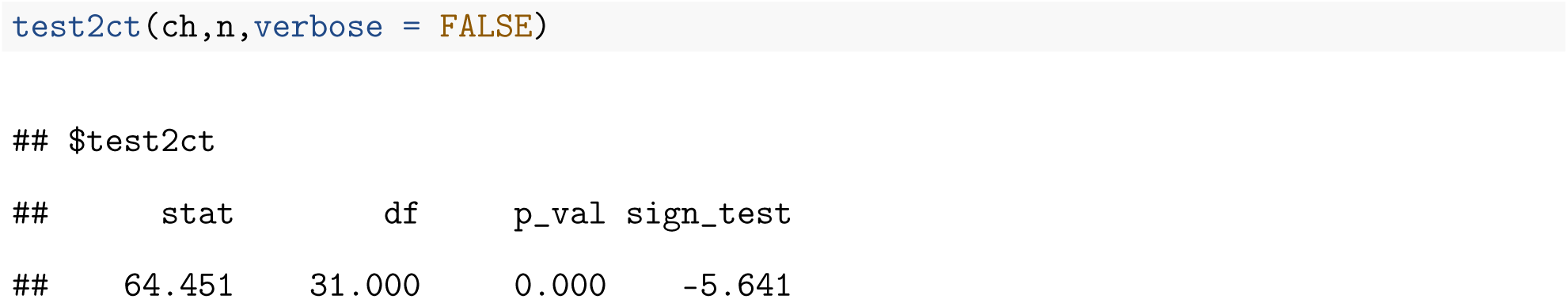

`Test 2.CT` is significant (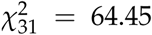, *P* < 0.01). We also provide the signed square root (`sign_test`) of the Pearson chi–square statistic as a directional test of the null hypothesis (Pradel et al. 2005), which is negative when there is an excess of individuals encountered at a given occasion among the individuals encountered at the previous occasion.

Note that, by default, the GOF functions in `R2ucare` returns all the contingency tables that compose the test under scrutiny, which might not be of immediate use and rather cumbersome on screen, hence the use of `verbose=FALSE` in the call to the `test2ct` function above. Now we ask whether there is a transient effect:

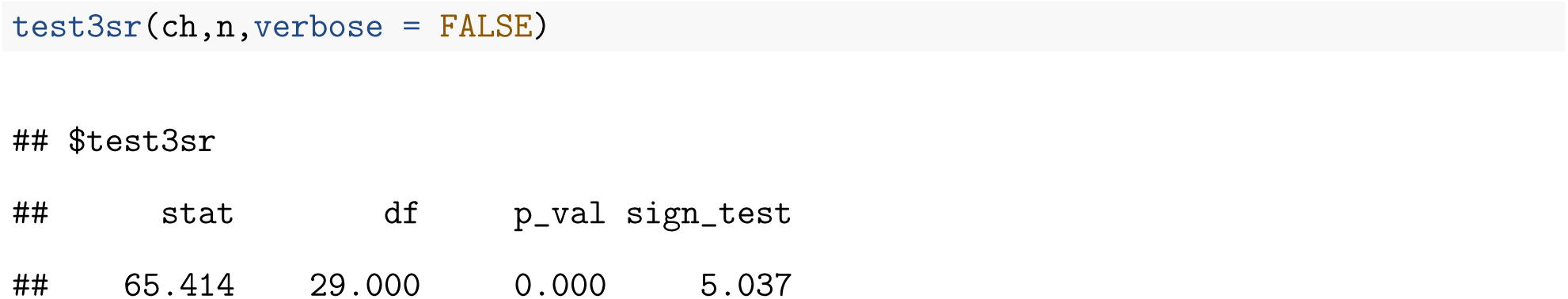

`Test 3.SR` is also significant (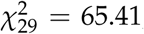, *P* < 0.01). We also provide the signed square root (`sign_test`) of the Pearson chi–square statistic (Pradel et al. 2005), which is positive when there is an excess of never seen again among the newly marked.

Navigating through the decision tree in Figure 1 suggests we should perform the two remaining tests:

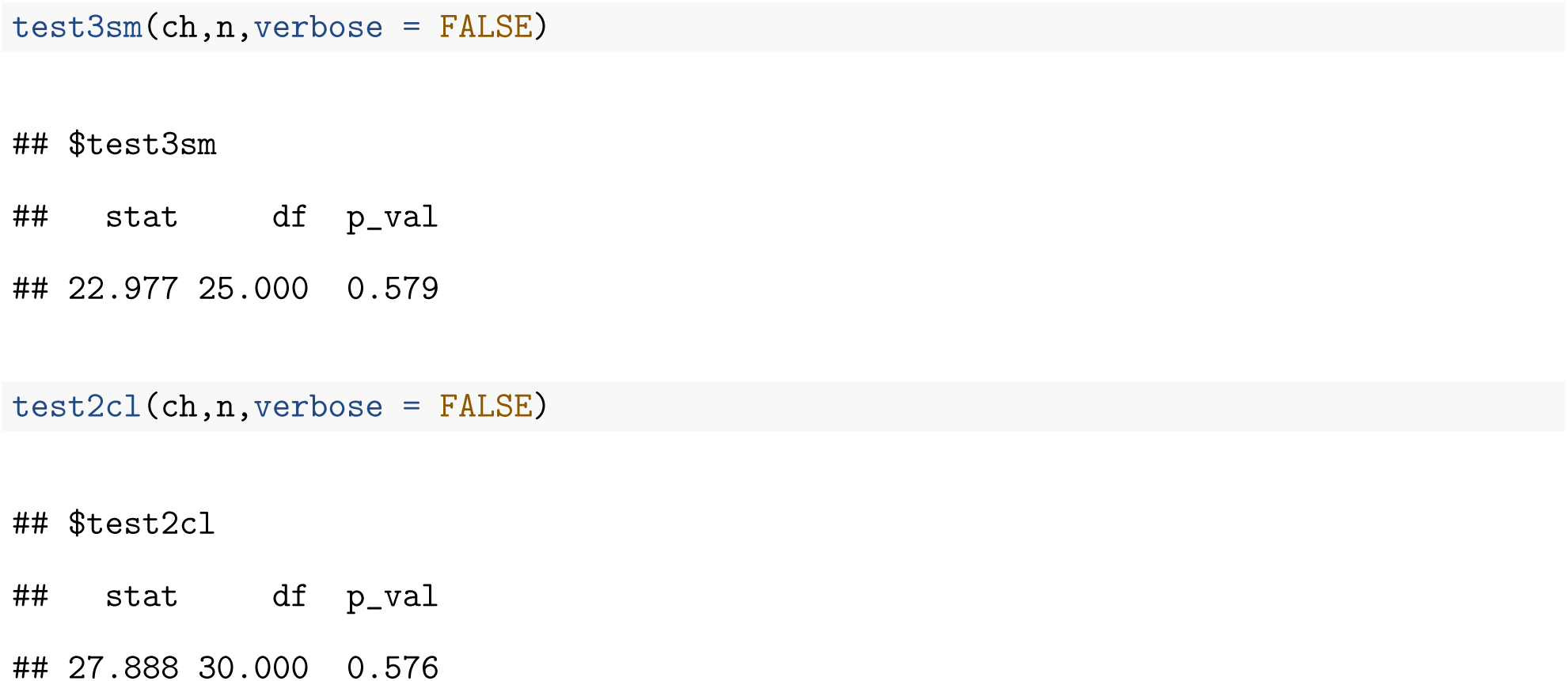

Neither `Test 3.Sm` (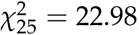, *P* = 0.58) nor `Test 2.CL` (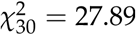, *P* = 0.58) is significant, therefore we recommend fitting a CJS model incorporating both a transience effect and a trap-dependence effect and start the analysis from there. In passing, it is possible to calculate a GOF test for this new model by removing the two components `Test 3.SR` and `Test 2.CT` to the overall GOF test (Pradel et al. 2005):

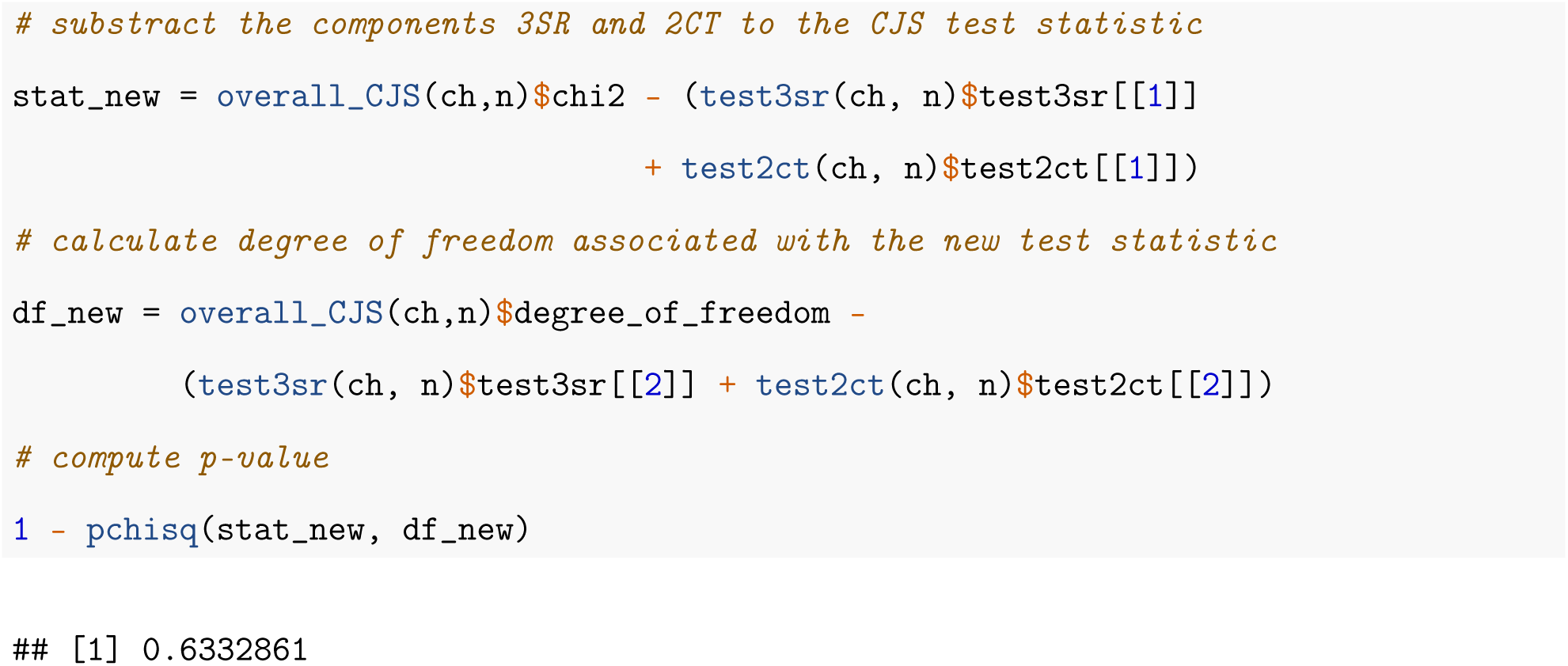

This new model incorporating transient and trap-dependence effects fits the wolf data well (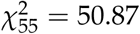, *P* = 0.63).

To date, no GOF test exists for models with individual covariates (unless we discretize them and use groups), individual time-varying covariates (unless we treat them as states) or temporal covariates; therefore, these covariates should be removed from the dataset before using `R2ucare`. For groups, we recommend treating the groups separately (see e.g. the example in the help file for `overall_CJS`).

## Goodness-of-fit tests for the Arnason-Schwarz model

We now wish to assess the GOF of the Arnason-Schwarz model to a dataset on Canada Geese (*Branta canadensis*) (Pradel et al. 2005). Briefly, the data consist of capture histories for 28,849 individuals marked and re–observed at wintering locations in the US between 1984 and 1986. We first read in the geese data that are provided with the package:

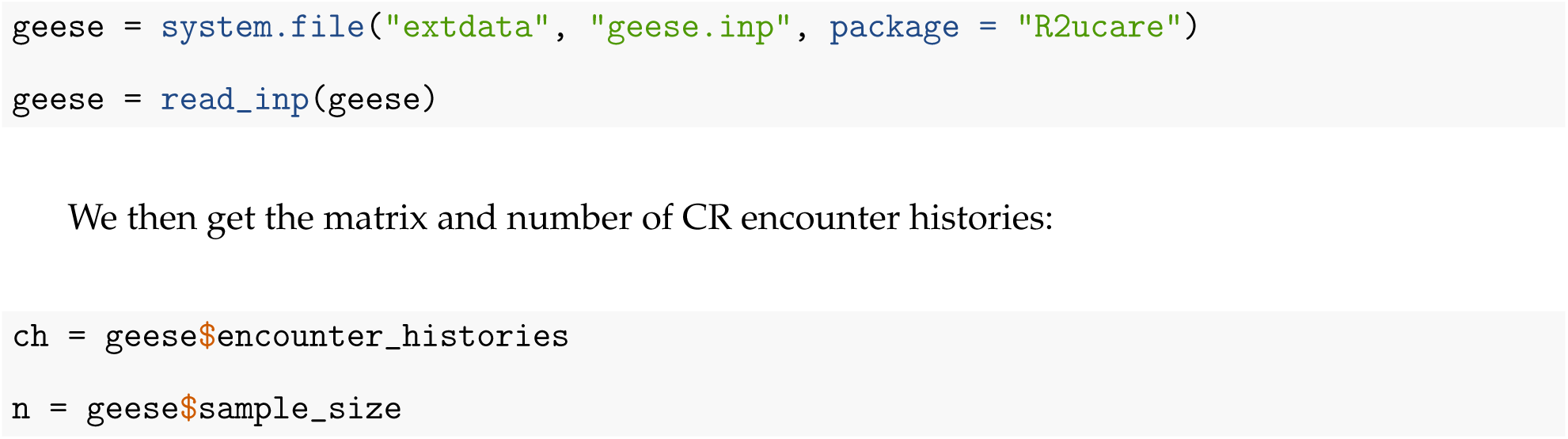

Then we assess the quality of fit of the Arnason-Schwarz model to the geese CR data with the `overall_JMV` function. Beware that it takes a minute or so to run the test because an iterative optimization procedure is involved to perform `Test M.ITEC` and `Test M.LTEC` (Pradel et al. 2003) that is repeated several times to try and avoid local minima.

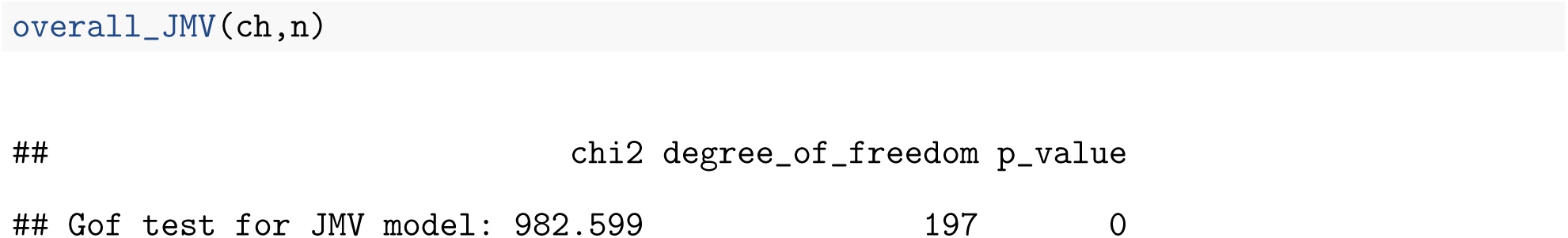

The null hypothesis that the Arnason-Schwarz provides an adequate fit to the data is clearly rejected (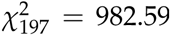, *P* < 0.01). In a second step, we further explore each component of the overall test:

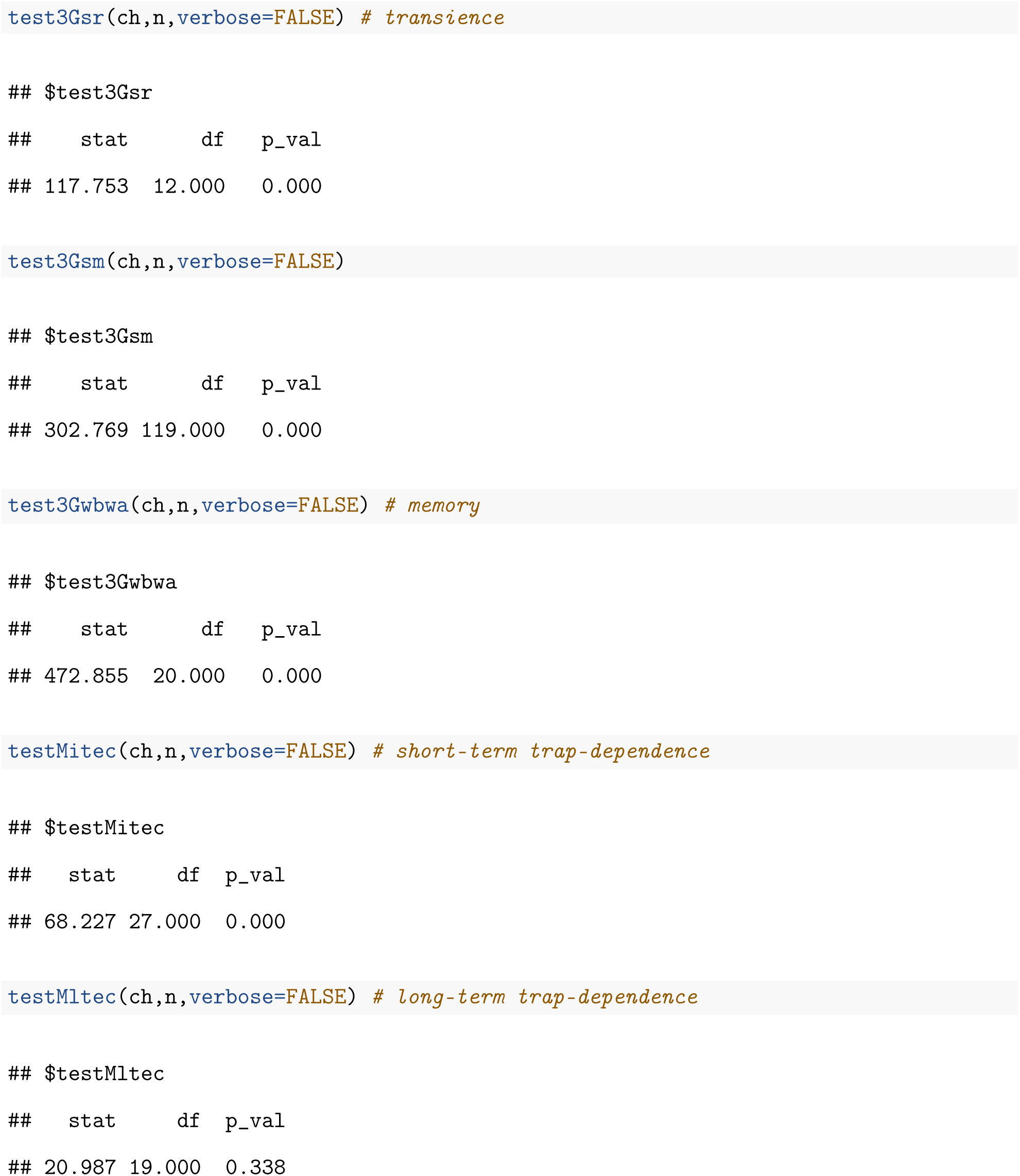

It appears that all components are significant but the test for a long-term trap-dependence effect. By setting the verbose argument to `TRUE` (by default argument), one could closely examine the individual contingency tables and better understand the reasons for the departure to the null hypotheses. For example, let us redo the test for transience `Test 3.GSR`:

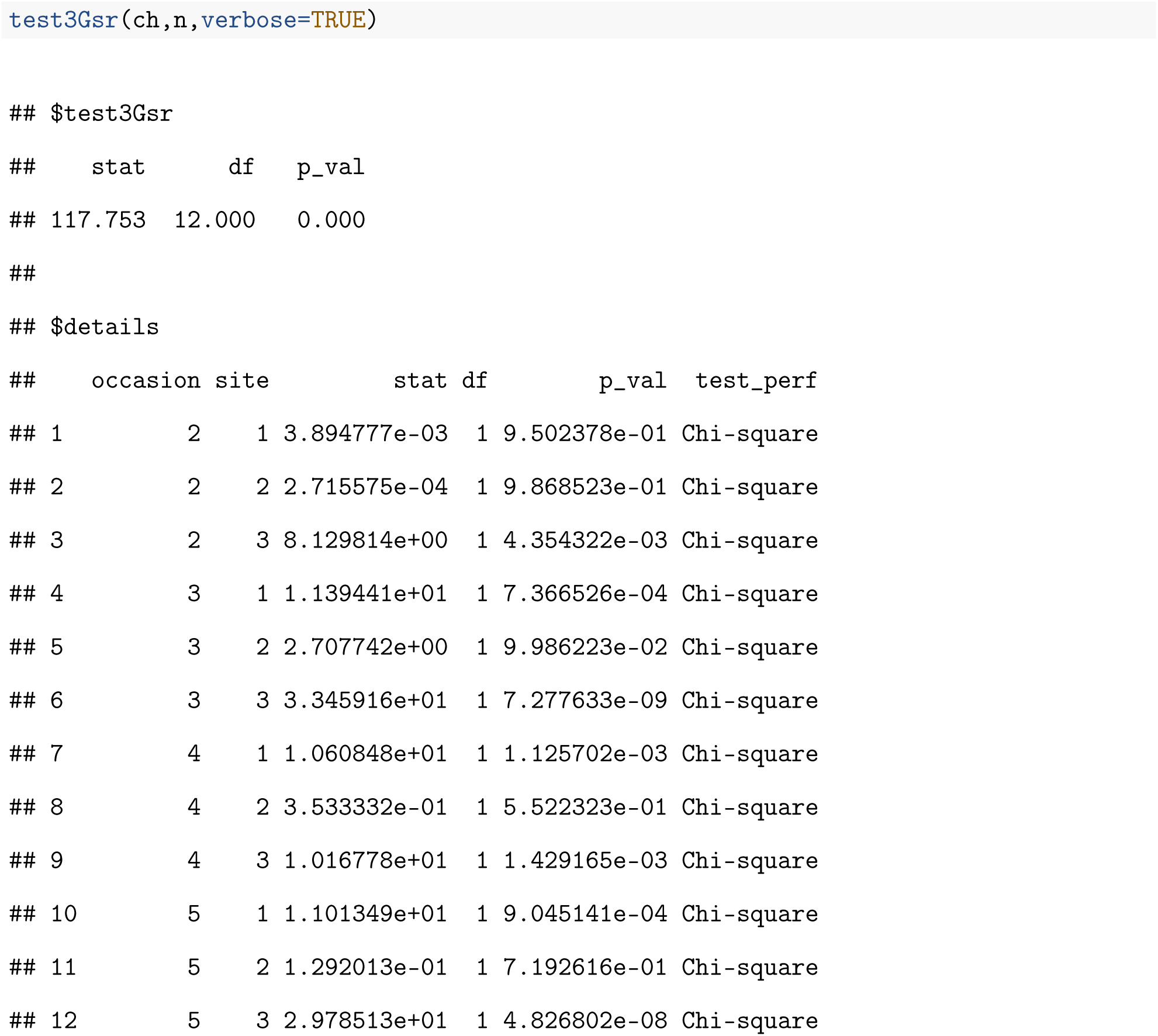

By inspecting the data.frame containing the details of the test, we see that there is no transients in site 2.

## Future directions

`R2ucare` allows evaluating the quality of fit of standard capture-recapture models for open populations. Future developments will focus on implementing goodness-of-fit tests for models combining different sources of data (McCrea et al. 2014) and residual-based diagnostics (Choquet et al. 2013, Warton et al. 2017).

## Availability

The current stable version of the package requires R 3.4.3 and is distributed under the GPL license. It can be installed from CRAN and loaded into a R session as follows:

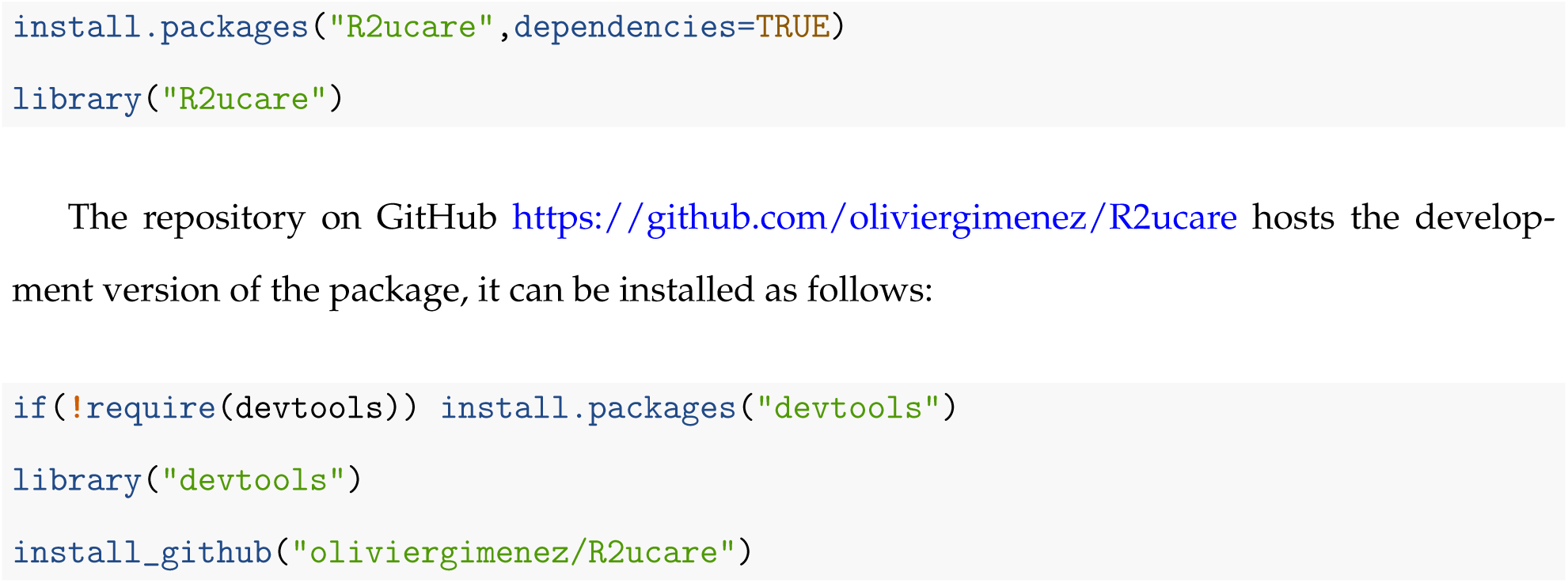

We also maintaina forumathttps://groups.google.com/forum/#!forum/esurge_ucare to which questions can be asked.

## Acknowledgments

Replication files (paper and code) are available on the first author’s Github account (https://github.com/oliviergimenez). This work was supported by a grant from the French National Research Agency, reference ANR-16-CE02-0007. We warmly thank E. Marboutin and J. Hestbeck for sharing the wolf and geese datasets, respectively.

## Authors’ contributions

OG, JDL and RP conceived the ideas and designed methodology; OG, JDL, RC and RP wrote the code; OG and RP led the writing of the manuscript. All authors contributed critically to the drafts and gave final approval for publication.

